# Dense poly(ethylene glycol) coatings maximize nanoparticle transport across lymphatic endothelial cells and accumulate in the skindraining lymph nodes

**DOI:** 10.1101/2020.08.01.232249

**Authors:** Jacob McCright, Colin Skeen, Jenny Yarmovsky, Katharina Maisel

## Abstract

Lymphatic vessels have recently been shown to effectively deliver immune modulatory therapies to the lymph nodes, which enhances their therapeutic efficacy. Prior work has shown that lymphatics transport 10–250 nm nanoparticles from peripheral tissues to the lymph node. However, the surface chemistry required to maximize this transport is poorly understood. Here, we determined the effect of surface poly(ethylene glycol) (PEG) density and size on nanoparticle transport across lymphatic endothelial cells (LECs) by differentially PEGylated model polystyrene nanoparticles. Using an established *in-vitro* lymphatic transport model, we found PEGylation improved the transport of 100 and 40 nm nanoparticles across LECs 50-fold compared to the unmodified nanoparticles and that transport is maximized when the PEG is in a dense brush conformation or high grafting density (R_f_/D = 4.9). We also determined that these trends are not size-dependent. PEGylating 40 nm nanoparticles improved transport efficiency across LECs 68-fold compared to unmodified nanoparticles. We also found that PEGylated 100 nm and 40 nm nanoparticles accumulate in lymph nodes within 4 hours after intradermal injection, while unmodified nanoparticles accumulated minimally. Dense PEGylation also led nanoparticles to travel the furthest distance from the injection site. Finally, we determined that nanoparticles are transported via both paracellular and transcellular mechanisms, and that PEG conformation modulates the cellular transport mechanisms. Our results suggest that PEG conformation is crucial to maximize nanoparticle transport across LECs and into lymphatic vessels, making PEG density a crucial design. Optimizing PEG density on nanoparticle formulations has the potential to enhance immunotherapeutic and vaccine outcomes.

## Introduction

Lymphatic vessels exist throughout the entire body and are known for transporting cells, fluid, and particulates from peripheral tissues to the local draining lymph nodes (LNs), where the adaptive immune response is formed [1]. In recent years, lymphatics have received increasing attention as potential drug delivery targets to transport immune modulatory therapies to the LNs without requiring direct injections. Delivering immunotherapies, including vaccines, to the LNs has been shown to potentiate their therapeutic effects, particularly crucial as efficacy of many immunotherapies still requires improvement. Recent studies have demonstrated that nanoparticles between 10 - 250 nm in diameter are transported preferentially via lymphatic vessels from peripheral tissues to LNs, highlighting that the transport functions of lymphatics can be taken advantage of for drug delivery [2-5].

While the size required for lymphatic entry is well established, conflicting data about the nanoparticle surface chemistry required to maximize lymphatic transport exist. Early studies on how size affects nanoparticle delivery to LNs demonstrated that 20 nm poly(ethylene glycol) (PEG)-stabilized nanoparticles were able to reach, and remain within, lymph node-resident dendritic cells compared to larger 100 nm nanoparticles [2]. Another study on the effects of nanoparticle size found that large 500 - 2000 nm, virus-like nanoparticles were taken up by skin-resident dendritic cells following intradermal injection, while smaller 50 - 200 nm virus-like nanoparticles were trafficked to the LNs via lymphatic drainage [3]. Combined, these initial studies provide evidence that the optimum nanoparticle size to reach LNs through lymphatic transport is within the 10 - 250 nm range.

Early studies demonstrated that coating nanoparticles with certain poloxamines, PEG-polypropylene oxide copolymers, can enhance LN accumulation of nanoparticles after intradermal administration. It was hypothesized that PEG chain length may contribute to some poloxamines enhancing nanoparticle transport more than others [6]. One study comparing cationic liposomes and anionic poly(lactic-co-glycolic acid) (PLGA) nanoparticles demonstrated that cationic liposomes accumulate to a greater extent in the LN after subcutaneous injection, compared to anionic PLGA nanoparticles [5]. However, there is a large size discrepancy between the two systems: cationic 180 nm liposomes were well within the lymphatic targeting size range, while anionic 350 nm PLGA nanoparticles were likely too large to preferentially enter lymphatic vessels. Another study demonstrated that positively charged 30 nm polyethyleneimine-stearic acid micelles preferentially accumulated in draining LNs compared to free antigen [7]. Another group demonstrated that coating 200 nm poly(methacrylate) nanoparticles with PEG markedly improved LN accumulation of nanoparticles after 12 and 48 hours [8]. Researchers also found that 50, 100, and 200 nm PEG-coated nanoparticles accumulated more in the LNs after subcutaneous injection compared to uncoated PLGA nanoparticles of the same size, suggesting that hydrophilicity is vital to maximize lymphatic transport of nanoparticles [9]. However, in this study, the surface potential of PEGylated nanoparticles was only −36.1 ± 14.6 mV, suggesting that the PEG coating was not very dense, as the methoxy-ended PEG would shield the negative charge of PLGA and reduce the nanoparticle surface potential. Similarly, researchers reported that PEGylation of poly-l-lysine dendrimers enhanced their transport to the LNs after subcutaneous injection. But it is unclear if the addition of PEG or the increase in size is primarily responsible for the improved LN accumulation, as dendrimers increased from 4 nm to 14 nm in diameter going from out of range to within range of size requirements for preferential transport by lymphatics [10]. Combined, these results suggest that hydrophilicity through addition of PEG, for example, may be beneficial for enhancing nanoparticle transport by lymphatics, but also highlight the importance to more critically assess the effect of surface chemistry, particularly PEG density, on nanoparticle transport by lymphatics.

PEGylating nanoparticles is a strategy that has been used extensively to enhance nanoparticle interactions with biological materials. PEGylation can improve nanoparticle drug delivery by reducing charge interactions with extracellular matrix and by preventing opsonization and phagocytosis by immune cells [11, 12]. Most notably, the chemotherapeutic drug doxorubicin, formulated as PEGylated liposomes, is one of the few FDA-approved nanoparticle treatments. The addition of PEG enhanced the circulation time of the chemotherapeutic, improving the overall efficacy [11-16]. PEG coatings can be optimized by modulating the density and molecular weight (MW) of PEG itself on the surface of nanoparticles. As PEG density on the nanoparticle surface is increased, the conformation of PEG transitions from a “mushroom” conformation that is more self-coiled to a “dense brush” conformation that is more linear, due to steric hindrances [17]. PEG conformation has been shown to be critical in enhancing nanoparticle transport across biological barriers. For example, to cross the mucus barrier, researchers have found that nanoparticles need to be in the “dense brush” conformation [18, 19]. Similarly, researchers have shown that only small (<100 nm) nanoparticles coated with PEG in the dense brush conformation effectively penetrate the interstitial tissue in the brain [20]. Additionally, tumor interstitial tissue penetration has also been improved by PEGylating nanoparticles: the addition of PEG to the surface of model, 60 nm negatively charged polystyrene (PS) nanoparticles (in an “intermediate brush” conformation) enhanced nanoparticle diffusion through breast cancer xenograft slices ex vivo compared to nanoparticles coated with PEG in the mushroom conformation [21]. These results demonstrate that the conformation of PEG on the surface is a key parameter that can affect nanoparticle delivery across biological barriers. However, how PEG density modulates nanoparticle transport by lymphatic vessels, and thus what the PEG density requirements are to maximize nanoparticle transport to the LNs and the mechanisms used by lymphatics to transport nanoparticles, remain poorly understood.

Here, we investigated the effect of PEG surface density on nanoparticle transport by lymphatic vessels and identified the PEG density required to maximize nanoparticle transport. We generated a library of nanoparticles coated with varying PEG densities, thus different PEG conformations, and tested their transport by lymphatic endothelial cells (LECs) using an established in vitro lymphatic transport model and in vivo after intradermal injection in mice [22]. Our resulting nanoparticle design criteria maximize nanoparticle transport to the LNs via lymphatic vessels, and therefore may enhance efficacy of immunotherapies and streamline the design of lymphatic targeting nanoparticle formulations.

## Materials and Methods

### Nanoparticle formulation

100 nm or 40 nm fluorescent carboxyl (COOH)-modified PS nanoparticles (Thermo Fisher Scientific, F8801) were covalently modified with 5 kDa MW methoxy-PEG-amine (NH_2_) (Creative PEGworks), as previously described [20]. Briefly, PS-COOH particles were suspended at 0.1% w/v in 200 mM borate buffer (pH = 8.2). Nanoparticles were generated with the following PEG concentrations: 350 μM (theoretical 100% PEG coverage of COOH groups), 175 μM (50% COOH groups), 87.5 μM (25% COOH groups), and 35 μM (10% COOH groups). PEG was conjugated to nanoparticles using 7 mM N-Hydroxysulfosuccinimide (NHS) (Sigma) and 0.02 mM 1-Ethyl-3-(3-dimethylaminopropyl) carbodiimide (EDC) (Invitrogen). The reaction was allowed to proceed on a rotary incubator at room temperature for at least 4 hours. Nanoparticles were collected using 100k MWCO centrifugal filters (Amicon Ultra; Millipore) and washed with deionized (DI) water. Nanoparticles were resuspended at 1% w/v in DI water and stored at 4°C.

### Nanoparticle characterization

Dynamic light scattering (DLS) was used to measure the hydrodynamic diameter and polydispersity index (PDI) of nanoparticles. Phase analysis light scattering (PALS) was used for measuring *ζ*-potential (NanoBrook Omni). Measurements were performed using a scattering angle of 90° at 25°C. Measurements were based on intensity of reflected light from scattered particles.

### PEG density characterization

PEG density was determined using a previously published method [23]. Briefly, 5kDa PEG-NH_2_ (Creative PEGworks) conjugated to fluorescein isothiocyanate (FITC) was conjugated to fluorescent (AlexaFluor®555) 100 nm carboxyl-modified nanoparticles. A FITC-PEG-NH_2_ standard curve was generated in DI water to calculate the PEG amount on the nanoparticle surface using a plate reader (Tecan Spark Multimode Microplate Reader). From these measurements, PEG grafting distance (D) and PEG density were estimated using the Flory radius of PEG (R_f_). The Flory radius of a polymer chain is defined as R_f_ ∼ αN^3/5^, where N is the degree of polymerization, and α is the effective monomer length. An unconstrained 5 kDa PEG chain has a R_f_ of 5.4 nm and occupies 22.7 nm^2^. PEG density and conformation can be correlated to the ratio of R_f_/D, with R_f_/D < 1-1.5 yielding a mushroom conformation, 1-1.5< R_f_/D > 4 yielding a brush conformation, and R_f_/D > 4 yielding a dense brush conformation.

Quantification of PEG on the surface of nanoparticles was performed using Fourier-Transform infrared (FTIR) spectroscopy (Vertex – 70 Bruker). PEGylated nanoparticle samples were scanned over a range of 400 – 4000 cm^-1^. The peak corresponding to the C-O-C ester linkages found in PEG chains was identified at 1083 cm^-1^ [24]. To quantify the amount of PEG on the surface of the nanoparticles, the intensity of the 1083 cm^-1^ peak was measured for known amounts of PEG, and a standard curve was generated. Using the same calculations as above, the R_f_/D value, which corresponds to the conformation of the PEG on the surface of a nanoparticle, was determined [24, 25].

### Nanoparticle uptake

Immortalized human LECs (hiLECs, [26]) were seeded at a density of 200,000 cells/cm^2^ onto collagen (Corning)-coated plates and cultured in endothelial growth media-2 (EGM2, Lonza) at 37°C and 5% CO_2_ overnight. hiLECs were incubated with 0.05% w/v nanoparticles for 3 h and uptake was assessed by flow cytometry or fluorescence microscopy. For fluorescence microscopy, samples were fixed with 2% paraformaldehyde (PFA, ThermoFisher) and imaged using a Zeiss Axio Observer. For flow cytometry, cells were released from the substrate using Accutase® (Innovative Cell Technologies), fixed with 2% PFA, and flow cytometry was performed using a BD FACSelecta. Data was analyzed using FlowJo software (Tree Star) and FIJI (ImageJ).

### Lymphatic transport model

Nanoparticle transport across LECs was assessed using an established in vitro model that recapitulates in vivo lymphatic transport [22]. Briefly, primary human dermal LECs (hLECs, Promocell C-12217) were seeded on 1.0 μm pore size, 12 mm transwell inserts (Falcon) at 200,000 cells/cm^2^ and cultured in EGM2 (Lonza) at 37°C and 5% CO_2_ for 48 h. Cells were pretreated with 1 μm/s transmural flow to simulate the tissue microenvironment. hLECs were treated with 1% w/v nanoparticles on the apical side and the basolateral compartment was sampled every 3 h for up to 24 h. Fluorescence intensity was measured using a plate reader (Tecan) and nanoparticles transported was calculated using a standard curve. Transport experiments were performed in EGM2 without growth factors to avoid the confounding effects of growth factors. To probe the transport mechanism the following transport inhibitors were used: 100 nM Adrenomedullin (Abcam ab276417), 62.5 μM Dynasore (Sigma D7693), or 62.5 μM Amiloride (Sigma A7410). Transport inhibitors were applied 2 hours prior to introduction of nanoparticles. Effective permeability was estimated using the following equation: 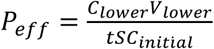, where C = concentration, V_lower_ = volume of the basolateral compartment, S = surface area, and t = time. hLEC monolayer integrity was confirmed after experiments using immunofluorescence.

### Immunofluorescence staining

Cells were fixed in 2% PFA for 15 minutes and incubated with mouse anti-human VE-Cadherin (BD Sciences) at 4°C overnight. Secondary antibodies conjugated to Alexa Fluor® 488 or 647 were used for detection (Thermo Fisher). Slides were mounted using DAPI (4’,6-diamidino-2-phenylindole)-containing Vectashield (Vector Laboratories Inc., Burlingame, CA) and imaged using a Zeiss Axio Observer. Image processing was performed using FIJI (NIH).

### C57Bl/6J lymphatic delivery model

10 μL of 5 mg/mL, fluorescently-labeled nanoparticles was intradermally administered to female C57Bl/6J mice (8-12 weeks old) in their forelimbs. Fluorescence intensity was measured using IVIS Spectrum Fluorescent & Chemiluminescent Imaging System (Caliper Life Sciences) over a 12h time period. Mice were anesthetized with isoflurane prior to nanoparticle injection and during imaging. Mice were euthanized after the final time point (8 or 12 h). Draining LNs were collected and homogenized to quantify the fluorescence signal from nanoparticles using a plate reader (Tecan). LNs were also fixed in 4% PFA for 6 hours and treated with a sucrose gradient. Tissues were then embedded within OCT (ThermoFisher), sectioned, and stained for FITC-B220 (BioLegend). Slides were mounted using DAPI (4’,6-diamidino-2-phenylindole)-containing Vectashield (Vector Laboratories Inc., Burlingame, CA) and imaged using a Zeiss Axio Observer. Image processing was performed using FIJI (NIH). All procedures were approved by the University of Maryland, College Park IACUC.

### Statistics

Group analysis was performed using a 2-way ANOVA, followed by Tukey’s post-test. Unpaired Student’s t-test was used to examine differences between only two groups. A value of *p* < 0.05 was considered significant (GraphPad). All data is presented as mean ± standard error of the mean (SEM).

## Results

### Increasing PEG density on nanoparticles neutralizes surface ζ-potential

The conformation of PEG on the surface of nanoparticles has been shown to affect how the nanoparticle interacts with surrounding tissues and cells [20, 23, 27]. In this study, we generated differentially PEGylated nanoparticles to determine how tuning PEG grafting density modulated surface PEG conformation and nanoparticle transport by LECs. We generated PS-COOH nanoparticles with varying PEG density, R_f_/D of 4.9 ± 0.1, 2.4 ± 0.1, 1.7 ± 0.1, and 1.3 ± 0.1 (**Fig 1A**). These R_f_/D values can be correlated to the conformation of PEG on the nanoparticle surface **(Fig 1B**). PEG grafting to the surface of the nanoparticle and R_f_/D values were confirmed using FTIR (**Fig 1C-D**). The 1083 cm^-1^ peak corresponds to C-O-C ester linkages characteristic of PEG chains. FTIR spectra of the additional nanoparticle formulations can be seen in the supplemental materials (**S1**). As expected, we found that increasing PEG density on nanoparticles slightly increased their diameter: unmodified PS-COOH nanoparticles had a diameter of 108 ± 1 nm, while addition of PEG increased nanoparticle diameter to 120 – 150 nm (**Fig 1E**). PEGylation also neutralized the negative surface charge of PS-COOH nanoparticles, from a ζ-potential of −22.4 ± 3.3 mV to −2.9 ± 2.5 mV (R_f_/D = 4.9), −5.1± 3.5 mV (R_f_/D = 2.4), −4.7 ± 2.5 mV (R_f_/D = 1.7), and −10.2 ± 6.6 mV (R_f_/D = 1.3) (**Fig 1F**). In addition, we confirmed stability of PEGylated nanoparticle formulations in EGM-2 media over 24 hours to ensure no aggregation occurred during the experimental time (**S2**).

**Figure 1:**
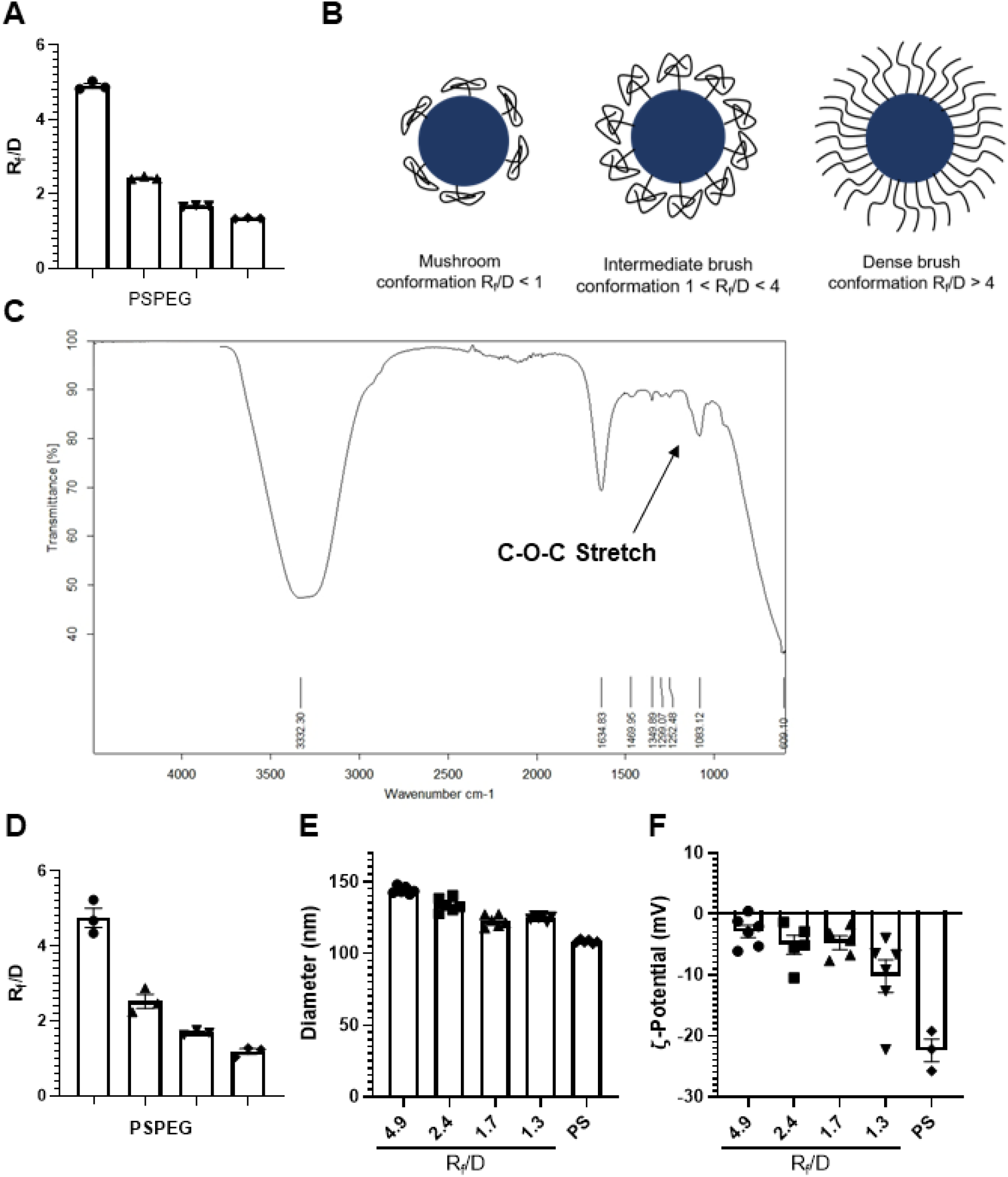
PEG grafted onto nanoparticles at different densities reduces surface charge. **A)** R_f_/D values of PSPEG measured via fluorescence. B) Schematic of PEG conformation on a model solid nanoparticle. **C)** Fourier Transform Infrared (FTIR) spectrum of PEG grafted on the surface of the nanoparticle. **D)** R_f_/D analysis as measured with FTIR. **E)** Dynamic Light Scattering (DLS) measurement of PEGylated NP diameter and **F)** Phase Analysis Light Scattering (PALS) measurement of NP ζ – potential. Data shown as mean ± SEM (n = 3 – 6).

### Dense brush PEG coatings on nanoparticles maximize their transport across LECs

We next sought to assess the effect of PEG density on nanoparticle transport by lymphatics. We used an in vitro transendothelial transport model (**Fig 2A**), where a monolayer of primary human LECs was cultured on the bottom of a collagen-coated transwell (**Fig 2B**) to simulate transport from the interstitium into the lymphatic vessel [22]. We found that the unmodified PS-COOH nanoparticles were minimally transported across LECs (0.03 ± 0.03%, **Fig 2B**) while 1.4 ± 0.3 % of densely PEGylated PSPEG_Rf/D = 4.9_ were transported after 6h (**Fig 2B**). By 24h there was a ∼90-fold increase in transport, with 4.2 ± 0.7% PSPEG_Rf/D = 4.9_ vs 0.05 ± 0.05% PS-COOH transported (**Fig 2A**). We found that the effective permeability (P_eff_) of the monolayer to nanoparticles also increased from 0.02 ± 0.02 μL/hr-cm^2^ for PS-COOH to 1.9 ± 0.3 μL/hr-cm^2^ for PSPEG_Rf/D = 4.9,_ without affecting the endothelial integrity (**Fig 2C-D**). Interestingly, any addition of PEG significantly increased nanoparticle transport by LECs after 24h (**Fig 2C**), but 1 < R_f_/D < 3 led to less transport than R_f_/D = 4.9 (**Fig 2C**). We also found that both PEGylated and unmodified nanoparticles were internalized by LECs (**Fig 2D**). To further probe if the effects of PEG density on nanoparticle transport across LECs translate across different nanoparticle sizes, we modified 40 nm PS-COOH nanoparticles with PEG (**S3**). We found that higher PEG density on 40 nm nanoparticles (R_f_/D = 4.4) enhanced nanoparticle transport across LECs compared to lower PEG density (R_f_/D = 0.9) (**Fig 2E**). Mass balance of nanoparticles in the top and bottom well were confirmed with >95% recovered after 24 hours (**S4**). These results indicate that the addition of PEG enhances the transport of nanoparticles across lymphatics and that “dense brush” PEG coatings maximize this transport.

**Figure 2:**
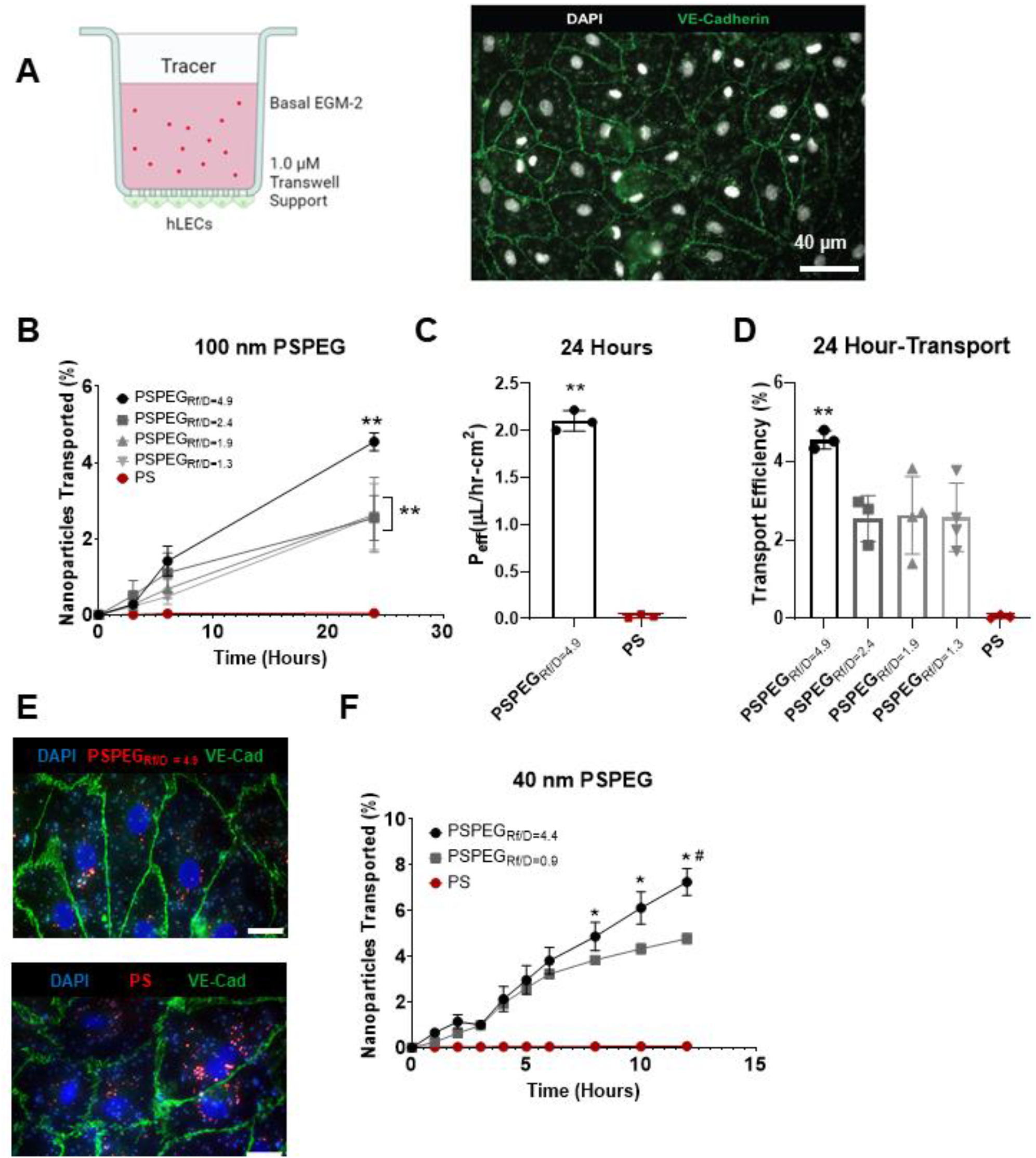
PEG Coating Improves Transport of 100 nm NP Across LECs. A) Schematic of transport model and representative image showing monolayer of LECs via VE-cadherin (green) and nuclei via DAPI (white). B) Representative images of LEC monolayer stained for VE-cadherin (green) and DAPI (blue) C) Percent of 100 nm NP transported across LEC monolayer over time. D) Measured effective permeability (P_eff_) of LEC monolayer to NP formulations. Percent of NP transported across LEC monolayer at the 24-hour time point. E) Representative images of LEC monolayer stained for VE-cadherin (green) and DAPI (blue) treated with unmodified PS or PSPEG NPs (red). (n = 3-4). F) Percent of 40 nm NP transported across LEC monolayer over time. Data presented as mean ± SEM (*p<0.05; **P<0.01; #P<0.01 comparing PSPEG_Rf/D=4.4_ and PSPEG_Rf/D=0.9_).

### PEG density on nanoparticles affects cellular mechanisms used by LECs to transport nanoparticles

To elucidate the mechanisms used by LECs to transport nanoparticles, we first investigated how PEG density on nanoparticles affects their uptake by LECs, as uptake is the first step in transcellular transport of materials. Using fluorescence microscopy and flow cytometry, we found that uptake of all nanoparticles by hiLECs was comparable, as indicated by the similar median fluorescence intensities (MFI) and similar number of cells positive for nanoparticles (> 85% for all nanoparticles, **Table 1**) at 3h, 6h, and 8h. Next, we investigated the cellular mechanisms involved in this transport using small molecule transport inhibitors to block the different cellular mechanisms. Several cellular mechanisms have been shown to be involved in nanoparticle transport across cellular barriers, including macropinocytosis, clathrin and/or caveolin-mediated endocytosis (micropinocytosis), as well as paracellular transport. We used amiloride to block macropinocytosis, adrenomedullin to reduce paracellular transport (by tightening cell-cell junctions, **Fig 3A**), and dynasore to inhibit the dynamin motor required for vesicle-based micropinocytosis. We found that 100 nm PSPEG_Rf/D = 4.9_ were not transported by macropinocytosis (**Fig 3B**). However, both adrenomedullin and Dynasore reduced 100 nm PSPEG_Rf/D = 4.9_ transport (**Fig 3B**), suggesting that both paracellular transport and micropinocytosis mechanisms are involved in nanoparticle transport by lymphatics. We also found that for the low PEG density, 100 nm PSPEG_Rf/D=1.3_ transport was reduced when each pathway was inhibited, suggesting paracellular transport, micropinocytosis, and macropinocytosis are all involved in transport of 100 nm PSPEG_Rf/D=1.3_ across LECs (**Fig 3C**).

**Table 1:** PEG conformation does not affect nanoparticle (NP) uptake in LECs. Median fluorescence intensity (MFI, imaging, n = 8) and % hiLECs positive for nanoparticles (flow cytometry, n = 3-4) after 3h., 6h, 8h.

| Incubation Time (h): |  | 3 h |  | 6 h |  | 8 h |  |
| --- | --- | --- | --- | --- | --- | --- | --- |
| NP type | Rf/D | MFI | % NP+ LECs | MFI | % NP+ LECs | MFI | % NP+ LECs |
| PSPEG | 4.9 ± 0.1 | 1.16 ± 0.36 | 91 ± 1 | 1.27 ± 0.28 | 92 ± 1 | 1.11 ± 0.18 | 94 ± 1 |
| PSPEG | 2.4 ± 0.1 | 0.93 ± 0.27 | 86 ± 1 | 1.02 ± 0.19 | 89 ± 2 | 1.05 ± 0.25 | 92 ± 2 |
| PSPEG | 1.7 ± 0.1 | 0.91 ± 0.23 | 91 ± 1 | 0.93 ± 0.31 | 90 ± 1 | 1.12 ± 0.35 | 92 ± 4 |
| PSPEG | 1.3 ± 0.1 | 0.93 ± 0.30 | 91 ± 1 | 1.05 ± 0.24 | 92 ± 1 | 0.98 ± 0.12 | 91 ± 2 |
| PS | - | 1.15 ± 0.23 | 91 ± 1 | 0.98 ± 0.17 | 91 ± 1 | 0.95 ± 0.27 | 89 ± 2 |

**Figure 3:**
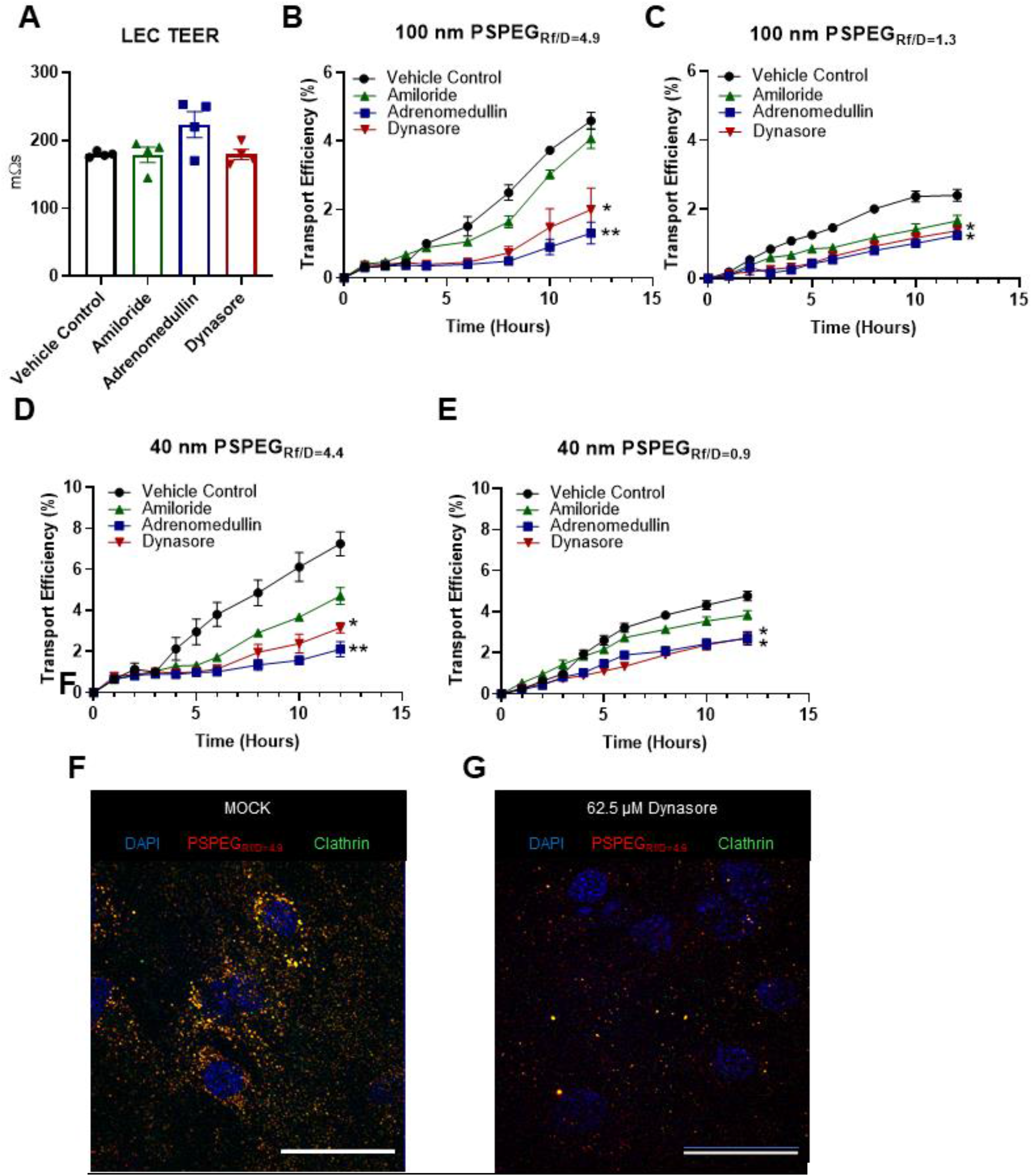
LEC *Both paracellular and transcellular transport mechanisms regulate Nanoparticle transport across LECs in-vitro*. **A)** Trans-endothelial electrical resistance (TEER) of the LEC monolayer after treatment with transport inhibitors. **B)** Transport efficiency of 100 nm PSPEG_Rf/D=4.9_ nanoparticle in the presence of transport inhibitors. **C)** Transport efficiency of 100 nm PSPEG_Rf/D=1.3_ NP in the presence of transport inhibitors. **D)** Transport efficiency of 40 nm PSPEG_Rf/D=4.4_ nanoparticle in the presence of transport inhibitors. **E)** Transport efficiency of 40 nm PSPEG_Rf/D=0.9_ nanoparticle in the presence of transport inhibitors. **F)** Confocal fluorescence image of PSPEG_Rf/D=4.9_ within LECs treated with the vehicle control and G) confocal fluorescence image of 100 nm PSPEG_Rf/D=4.9_ within LECs treated with Dynasore Scale bar: 30 μm. (n = 3-4) Data presented as mean ± SEM (*p<0.05; **P<0.01).

To further probe the mechanisms of transport, and the nanoparticle characteristics regulating lymphatic transport, we used differentially PEGylated 40 nm nanoparticles (**S2**). When 40 nm densely PEGylated nanoparticles (PSPEG_Rf/D=4.4_) were introduced to the transport model, we found that dynasore, adrenomedullin, and amiloride were each able to reduce transport (**Fig 3D**), suggesting that paracellular transport, micropinocytosis, and macropinocytosis are all involved in 40nm PSPEG_Rf/D=4.4_ transport across LECs. The transport of the 40 nm PSPEG_Rf/D=0.9_ was affected by transport inhibitors in a similar fashion as the 100 nm PSPEG_Rf/D = 4.9_, with adrenomedullin and Dynasore reducing transport, but amiloride having no significant effect, (**Fig 3E**), suggesting that only micropinocytosis and paracellular transport are involved. To further investigate the specific micropinocytosis mechanisms, we used fluorescent microscopy to determine if nanoparticles colocalized with known endocytosis mediators. We found that 100 nm PEGylated nanoparticles colocalized with clathrin in LECs, suggesting that clathrin-mediated endocytosis is one of the mechanisms involved in 100 nm PSPEG_Rf/D = 4.9_ transport by LECs (**Fig 3F, S5**). The mechanisms by which PEG density modulates nanoparticle transport mechanisms used by LECs are currently under investigation in our lab.

### Densely PEGylated nanoparticles accumulate in the LNs in vivo

To confirm that our in vitro findings are representative of in vivo lymphatic transport, we probed nanoparticle accumulation in the LNs over 12 h after intradermal injection in mice. We found that minimal amounts of 100 nm PS-COOH nanoparticles were transported to the LNs even after 12 h, while 100 nm PSPEG_Rf/D = 4.9_ nanoparticles were transported to the LNs as early as 4 h after injection (**Fig 4A**). We measured the distance from injection site and found that after 12 h, 100 nm PS-COOH beads traveled 0.24 ± 0.04 cm from the injection site, whereas 100 nm PSPEG_Rf/D = 1.3_ traveled 0.61 ± 0.01 cm from the injection site (**Fig 4B**). 100 nm PSPEG_Rf/D = 4.9_ traveled the furthest from the injection site, measuring 0.73 ± 0.04 cm from the nanoparticle injection site. Distance traveled was significantly higher for 100 nm PSPEG_Rf/D=4.9_ at 8 h and 12 h compared to both 100 nm PS-COOH and PSPEG_Rf/D=1.3_. Both PEGylated 100 nm nanoparticle formulations significantly improved distance traveled compared to unmodified 100 nm PS-COOH nanoparticles (**Fig 4B**), further indicating that a dense coating of PEG optimizes transport of nanoparticles to LNs. When we quantified fluorescence in draining LNs, we found that both100 nm PSPEG_Rf/D=4.9_ and PSPEG_Rf/D=1.3_ accumulated in the LN after 8h whereas 100 nm PS-COOH had a significantly reduced signal (**Fig 4C-F**). To examine the size dependence on in-vivo transport we intradermally administered densely PEGylated 40 nm nanoparticles (PSPEG_Rf/D=4.4_), sparsely PEGylated 40 nm nanoparticles (PSPEG_Rf/D=0.9_), and 40 nm PS-COOH nanoparticles thenmeasured transport to draining LNs via IVIS. Like the 100 nm formulations, PEGylated 40 nm nanoparticles began to appear in draining LNs 4 h after intradermal injection (**Fig 5A**). Similar to our findings with 100 nm nanoparticles, the addition of some PEG increased the distance 40 nm nanoparticles traveled from 0.29 ± 0.05 cm by 40 nm PS-COOH to 0.69 ± 0.09 cm by 40 nm PSPEG_Rf/D=0.9_. 40 nm PSPEG_Rf/D=4.4_ were found to travel furthest from the injection site, with 0.88 ± 0.06 cm after 12h (**Fig 5B**). When we quantified fluorescence in draining LNs, we found that both 40 nm PSPEG_Rf/D=4.4_ and PSPEG_Rf/D=0.9_ accumulated in the LN after 8h whereas 40 nm PS-COOH had a significantly lower signal (**Fig 5D-F**). We then examined nanoparticle localization within draining LNs using IF imaging. IF imaging confirmed IVIS results, with no PS-COOH signal observed in the LNs after 8h. PSPEG_Rf/D=4.9_ was observed within the LN after 8h for both 40 nm and 100 nm nanoparticle sizes. PEGylated formulations appear to be localized within the subcapsular sinus, suggesting that lymphatic drainage through the afferent lymphatic vessels mediated transport to the LN (**Fig 5C, 4C**).

**Figure 4:**
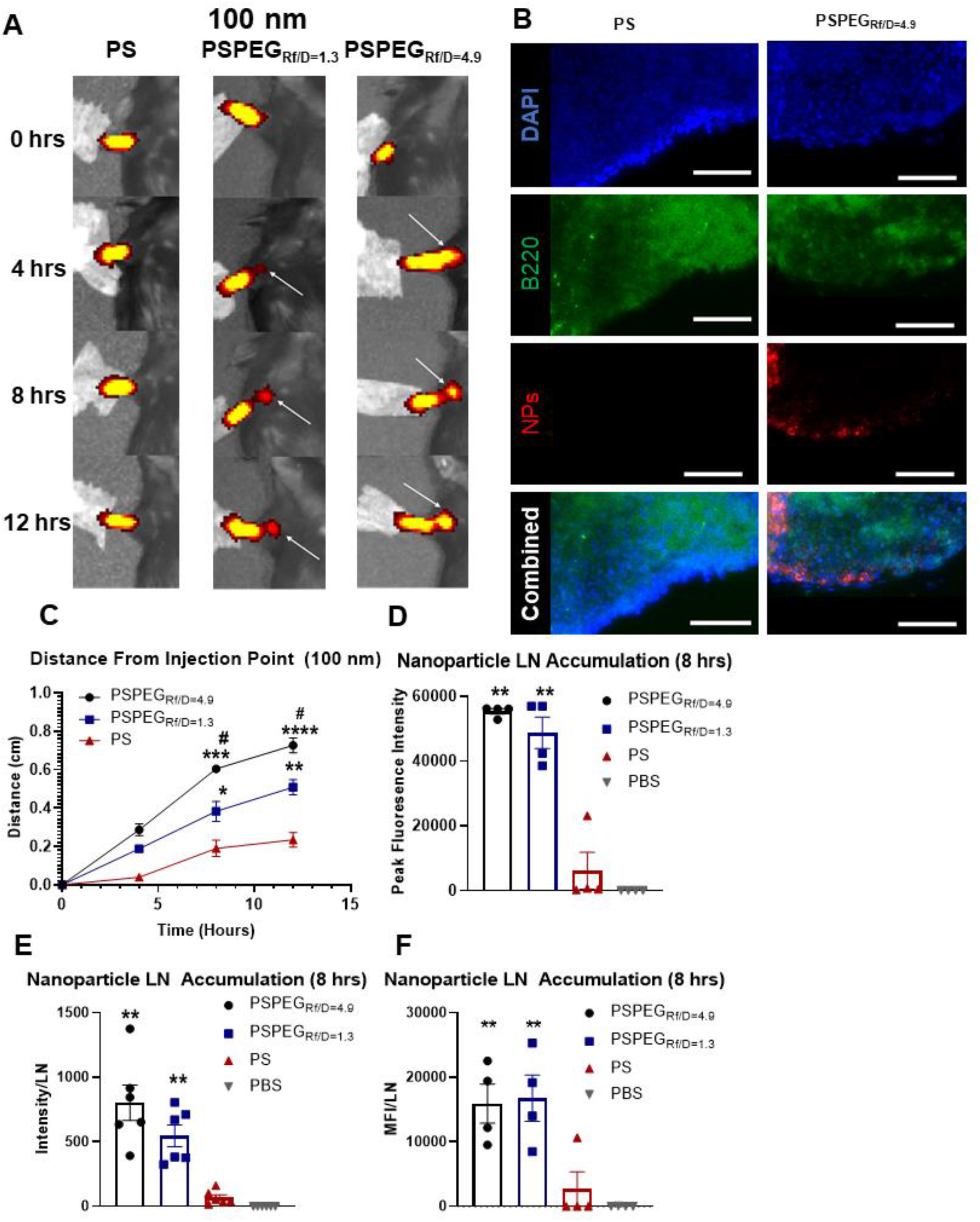
Dense-Brush PEG Coating Required for Improved Lymphatic Targeting In-Vivo. **A)** Representative images of intradermally injected 100nm NPs in C57Bl/6J mice up to 12h post injection measured using IVIS. White arrow indicates accumulation within lymph nodes. **B)** Nanoparticle transport measured as maximal distance of fluorescent signal from injection site. **C)** Lymph node sections stained for DAPI (nucleus) and B220 (B-cells) after injection of 100 nm PS or PSPEG_Rf/D=4.9_ nanoparticles. **D)** Nanoparticle accumulation within LN measured as average fluorescence signal over the area of dissected LN (MFI/LN) and as **E)** Peak fluorescence signal. **F)** Fluorescence signal of homogenized lymph nodes. (n = 6) Data presented as mean ± SEM (*p<0.05; **P<0.01, *** P<0.001; #P<0.01 comparing PSPEG_Rf/D=4.9_ and PSPEG_Rf/D=1.3_).

**Figure 5:**
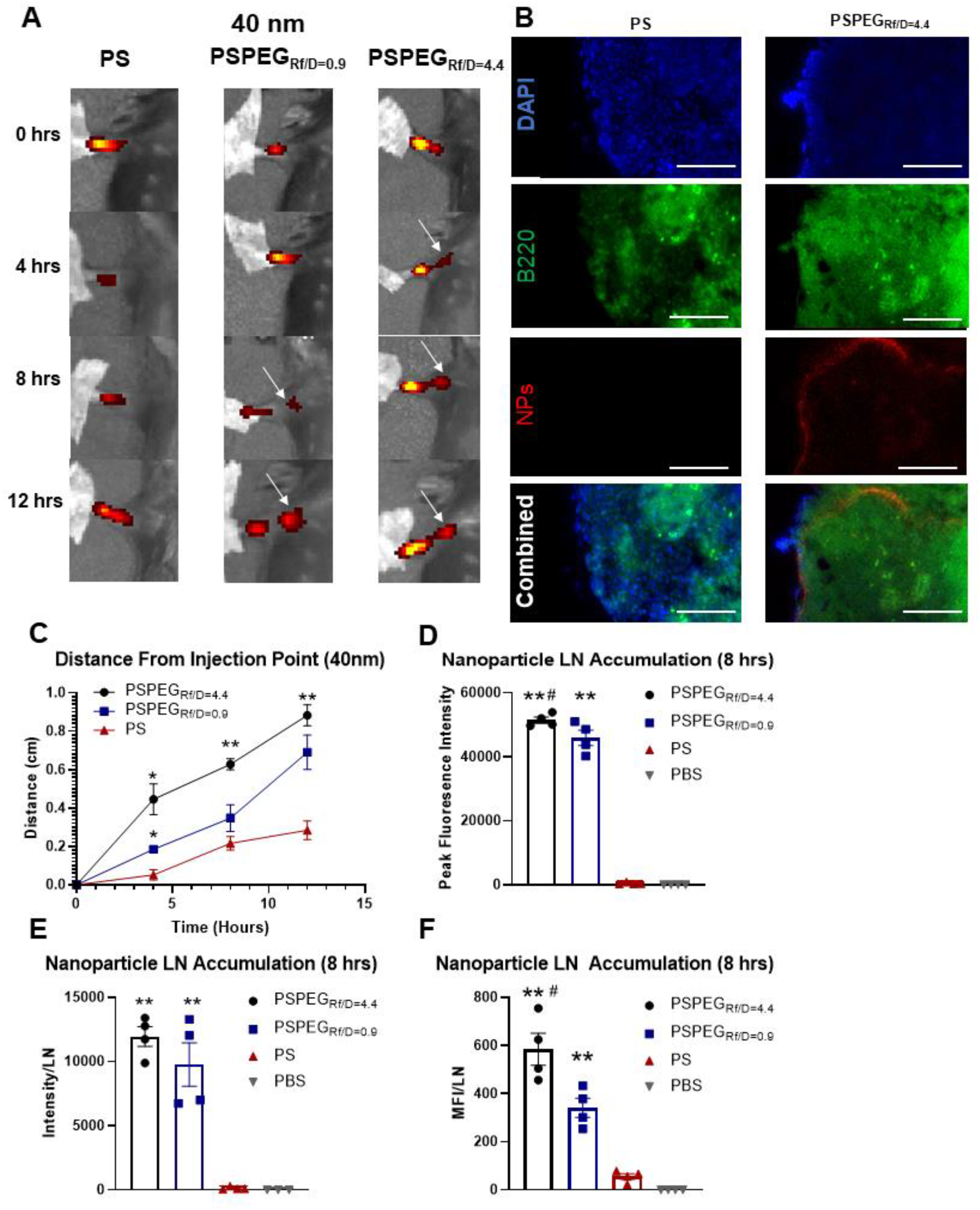
Dense-Brush PEG Coating Required for Improved Lymphatic Targeting In-Vivo. **A)** Representative images of fluorescent signal from intradermally injected 40 nm NPs in C57Bl/6J mice at different times post injection measured using IVIS. White arrow indicates accumulation within lymph nodes. **B)** Transport measured as maximal distance of fluorescent signal from injection site.. **C)** Lymph node sections stained for DAPI (nucleus) and B220 (B-cells) with nanoparticles seen in red for 40 nm PS and PSPEG_Rf/D=4.4_. **D)** Nanoparticle accumulation within LN measured as average fluorescence signal over the area of dissected LN (MFI/LN) and as **E)** peak fluorescence signal. **F)** Fluorescence signal of homogenized lymph nodes. (n = 6) Data presented as mean ± SEM (*p<0.05; **P<0.01; #P<0.01 comparing PSPEG_Rf/D=4.4_ and PSPEG_Rf/D=0.9_).

## Discussion

In this work, we investigated the effects of PEG surface density on nanoparticle transport across lymphatics. We found that in vitro, addition of any amount of PEG to nanoparticle surfaces increased their transport across lymphatics compared to hydrophobic nanoparticles, and a “dense brush” of PEG (R_f_/D > 4) on nanoparticles maximized this transport. We further found that densely PEGylated nanoparticles were more effectively transported to skin-draining LNs after intradermal injection compared to nanoparticles with lower or no PEG coating. Finally, we found that macropinocytosis, micropinocytosis, and paracellular transport mechanisms mediated transport of PEGylated nanoparticles across lymphatics, but that macropinocytosis did not occur in all PEG conformations. In summary, we identified that “dense brush” PEG coatings maximize nanoparticle transport into the lymphatics and thus the down-stream LNs.

PEG has been used to modulate nanoparticle transport across endothelial barriers, including blood endothelium, lymphatic endothelium, and the blood-brain barrier. Generally, PEG has been shown to enhance systemic circulation and blood endothelium needs to be targeted using additional vectors such as ICAM1 or sugars that target specific ligands on the blood endothelium [28, 29]. This is likely due to the high flow rates in blood vessels that cause nanoparticles not to come into close contact with endothelial cells for long enough to allow uptake and transport across the endothelium [30, 31]. A recent study used 100 nm poly(lactic acid)-PEG nanoparticles with surface PEG ranging from 1 – 10 kDa to probe the effects of PEG MW on transcytosis across brain vascular endothelial monolayers [32]. PEG density on the surface of the nanoparticles was maintained at 17-20 PEG/100nm^2^. It was found that the higher molecular weight PEG polymers displayed improved transport across the monolayers, with 60% translocation efficiency of nanoparticles coated with 5 kDa and 10 kDa MW PEG and only 20% translocation efficiency of nanoparticles coated with 1 kDa PEG. Interestingly, the maintained PEG density translates to R_f_/D > 2 for nanoparticles coated with 5 kDa and 10 kDa, while R_f_/D < 1 for 1 kDa coated nanoparticles, indicating that PEG on 1kDa nanoparticles was in mushroom conformation, while PEG was in brush conformation for 5 kDa and 10 kDa nanoparticles. Another study by Rabanel et al suggested that 5kDa PEG coatings increased uptake of nanoparticles into brain endothelial cells, but PEG MW did not have any effect on transport across them [33]. Kim et al demonstrated that PEGylating ionizable lipid nanoparticles reduced their uptake in the liver [34]. They hypothesized that this was due to reduced ApoE protein on the nanoparticle surface, which is one of the key mechanisms that leads to nanoparticle uptake in the liver. Additionally, work by Williams et al showed that PEG enhanced kidney accumulation of nanoparticles and that this was likely due to endocytosis of nanoparticles by the peritubular endothelium [35]. Our findings showed that PEGylation enhances nanoparticle transport across LECs, suggesting that the type of endothelium and other factors such as contact time can affect whether PEG improves transendothelial transport of nanoparticles. For transport into lymphatic vessels, our findings that PEG enhances nanoparticle transport across lymphatics are corroborated by prior work indicating that nanoparticles and liposomes with PEG coatings (with undefined PEG densities) transport effectively to the LNs. Our work adds an additional layer of understanding that a high PEG density maximizes this transport [5-10].

Surface PEG conformation on nanoparticles has been shown to affect nanoparticle uptake by cells [36]. Several studies have shown that as PEG density increases on nanoparticles, uptake by macrophages and dendritic cells [37-42], as well as cancer cells [37, 43], is reduced. PEGylation appears to have differing results depending on the cell type – PEGylation reduces nanoparticle uptake by macrophages and dendritic cells, as indicated by studies demonstrating that PEG needed to be in a brush conformation to evade uptake and clearance by macrophages [41]. However, addition of PEG to the surface of nanoparticles increased their uptake by neutrophils [23] and several cancer cell types (HeLA, MDA-MB231, VK2) [37, 44]. Interestingly, several studies suggest that changes in protein corona on nanoparticle surface with changing PEG density may in part be responsible for modulating cellular uptake [36, 37, 39, 43]. One study demonstrated that without a protein corona, some PEG, but not high density/MW PEG, could increase uptake by prostate cancer cells compared to no PEG, and without protein, no PEG was optimal [43]. Another study demonstrated that a double layer of PEG, with a second layer having a mushroom conformation (low density of PEG, R_f_/D < 1.5) reduced protein binding affinity but not total protein binding on the nanoparticle surface. They found that this second layer of PEG reduced nanoparticle uptake by liver sinusoidal endothelial cells [36]. In our studies, we found that PEG density did not affect nanoparticle uptake by LECs, and that it only modulated nanoparticle transport across LECs (**Table 1**). The non-phagocytic nature of LECs may account for some of these differences. Additionally, we are currently investigating effects of the protein corona on nanoparticle transport across lymphatics, as changes in protein corona appear to be strongly linked with differential uptake and transport of nanoparticles.

Nanoparticle transport across biological barriers, like the endothelium, has been shown to be governed by macro- and micropinocytosis, as well as paracellular transport mechanisms. Studies have shown that larger nanoparticles (>200 nm in size) are transported across cellular barriers via macropinocytosis, while smaller nanoparticles are often transported by various mechanisms of micropinocytosis. Here, we found that densely PEGylated 130 nm nanoparticles are transported via micropinocytosis, likely clathrin-mediated endocytosis, and paracellular transport routes across LECs. This is consistent with prior studies showing that albumin, a 10 nm globular nanoparticle-like protein, is transported across lymphatics via both clathrin- and caveolin-mediated, as well as paracellular transport mechanisms. Additionally, researchers have demonstrated that macro- and micropinocytosis are involved in transport across endothelial cells in tumors and the blood brain barrier [45-51]. Rabanel et al demonstrated that nanoparticles coated with 5 kDa PEG were taken up primarily via macropinocytosis pathways in brain endothelial cells [33]. Tehrani et al found that inhibiting micropinocytosis reduced transcytosis across brain endothelial cells by 60% for 5 kDa PEG-coated nanoparticles, while transcytosis of 2 kDa PEG-coated nanoparticles was reduced only by 25% after inhibiting micropinocytosis [32]. These findings suggest that nanoparticle uptake and transcytosis pathways may differ with different MW and density of PEG, corroborating our findings. A study aiming to improve the circulating time of nanoparticles found that maintaining PEG surface conformation within the intermediate brush domain prevented non-Kupffer cell uptake in the liver. They observed that these intermediately PEGylated nanoparticles were the least preferentially taken up by endothelial cells, accounting for the improved circulation times, and the densely PEGylated nanoparticles were taken up at higher rates [36]. Additionally, studies have demonstrated that nanoparticle transport across brain microvascular endothelium can be enhanced by taking advantage of existing receptor-mediated transcytosis, such as that of albumin (clathrin/caveolin-dependent) [49, 52-55]. Altogether, a variety of factors appear to influence transendothelial transport mechanisms, and data from the literature and our study suggest that there may be differences in mechanisms depending on the tissue, type of endothelium, and pathological condition [51].

For nanoparticles to reach cells within a tissue, including lymphatic vessels, they need to cross the extracellular matrix (ECM). The ECM forms a hydrogel-like structure composed of fluid, solutes, fibrillar proteins, and proteoglycans, such as collagen. Studies on nanoparticle-ECM interactions have demonstrated that size, shape, and charge are key considerations when designing nanoparticle therapies that need to cross ECM barriers. Large nanoparticles are restricted from crossing the ECM barrier through steric hindrance via the mesh spacing produced by fibers within the ECM. This spacing can become extremely restrictive within tissues. For example, the basement membrane mesh of the subcapsular sinus of the LN forms a tight mesh that prevents molecules larger than 70 kDa from entering the LN via afferent lymphatic vessels [56]. This restrictive barrier can be observed in action in our study here, where IF images of LN slices show nanoparticles sequestered on the edge of the LN, within the subcapsular sinus. Restrictive extracellular spacing can be seen in many other tissues, with the spacing often estimated between 20-60 nm in diameter, suggesting that tissues are impermeable to >100 nm sized particles [57]. However, this notion has been challenged recently: Nance et al have demonstrated that particles as large as 114 nm in diameter were able to diffuse within human and rat brain tissue. Indeed, their study highlighted that the pore size within in-vivo ECM is highly heterogenous, with them concluding that within the brain tissue samples, more than one quarter of all pores were >100 nm in diameter [20]. Our studies similarly suggest that skin ECM may be permeable to nanoparticles up to 150 nm in diameter.

Surface chemistry is another key factor affecting nanoparticle transport across ECM barriers. Several studies have demonstrated that nanoparticles with charge opposing that of the fibrous materials of a hydrogel, like ECM, have reduced diffusion within the gel or ECM space [58-62]. Additionally, some studies suggest that repulsive charges can enhance nanoparticle diffusion across ECM barriers compared to attractive charges, but this effect may be minimal [58-62]. Most studies indicate that neutral charge leads to the highest diffusion of nanoparticles across ECM [20, 60, 62], suggesting that PEGylating nanoparticles as we have done in our studies also enhances their transport across ECM. However, fewer studies have assessed how PEG density affects nanoparticle transport across the ECM. One study found that increased PEG density can enhance nanoparticle diffusion through collagen-based ECM materials [63], but this was not recapitulated in matrigel-based gels. Furthermore in this study, PEG density changes are a result of using PEG chains with different molecular weights (MW), which could also affect diffusion, e.g., due to entanglement with ECM chains for larger MWs. Another study has demonstrated that increasing PEG density enhanced liposome diffusion in collagen-based ECM hydrogels [64], and similarly previous work has shown that dense PEG coatings enhanced nanoparticle diffusion through brain ECM [20]. In our studies here, we show that PEG density modulates nanoparticle transport to the LNs, which may be due to both reduced transport across lymphatic endothelial cells as well as ECM, based on these existing studies, and is a current topic of investigation in our lab.

In summary, our study demonstrates that the addition of PEG to the surface of hydrophobic nanoparticles, particularly as dense PEG coatings (R_f_/D >4), enhances nanoparticle transport into lymphatic vessels. Our results are consistent with prior work demonstrating that PEG enhances uptake and transport across other endothelial barriers as well as the cellular mechanisms involved in this transport. Our study is the first to directly correlate PEG density and efficiency of lymphatic transport of nanoparticles. Additionally, it is the first to demonstrate that densely PEGylated nanoparticles are transported via both micropinocytosis and paracellular transport mechanisms, but not macropinocytosis, by LECs. Furthermore, we demonstrated that PEG density and size can affect the cellular mechanisms used to transport nanoparticles across lymphatic barriers. Thus, our work highlights that dense PEG coatings are the optimal strategy to formulate nanoparticles that maximize transport across lymphatics and to the LNs. Our work also sheds new light onto the effects of PEG density on cellular mechanisms of nanoparticle transport. Our findings are particularly crucial for future development of immune modulatory therapeutic strategies.

## Abbreviations

LN: lymph node
PEG: poly(ethylene glycol
PS: Polystyrene
PLGA: poly(lactic-co-glycolic acid
DI: Deionized
DLS: dynamic light scattering
PDI: polydispersity index
PALS: phase analysis light scattering
R_f_: Flory radius
D: grafting distance
LECs: lymphatic endothelial cells

## Supplemental Figures

**Supplemental Figure 1:**
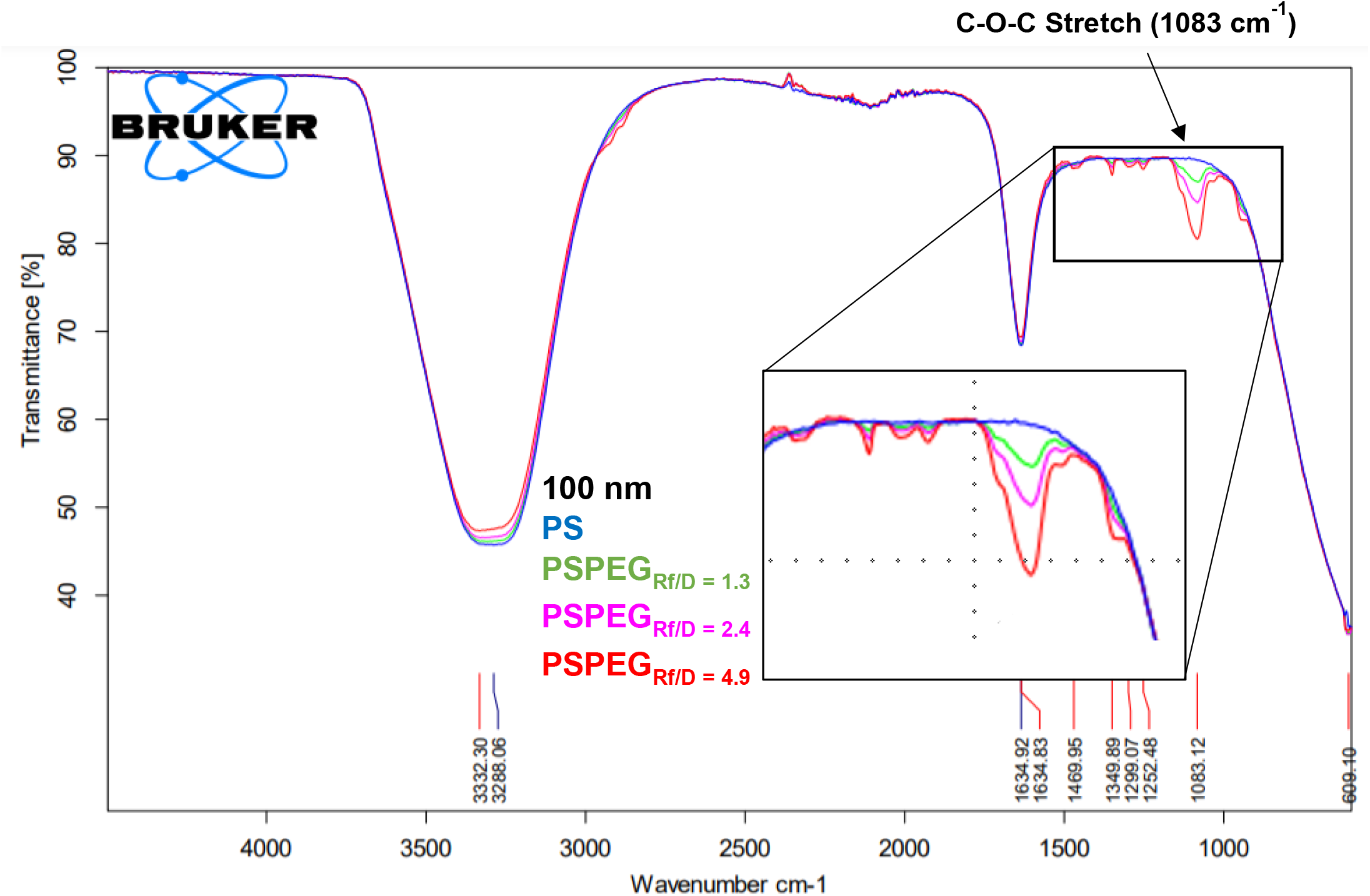
FTIR Spectra of 100 nm PSPEG Formulations: Representative spectra of PS, PSPEG_Rf/D = 1.3,_ PSPEG_Rf/D = 2.4_, and PSPEG_Rf/D = 4.9_ with ester linkage indicated at 1083 cm-1 peak.

**Supplemental Image 2:**
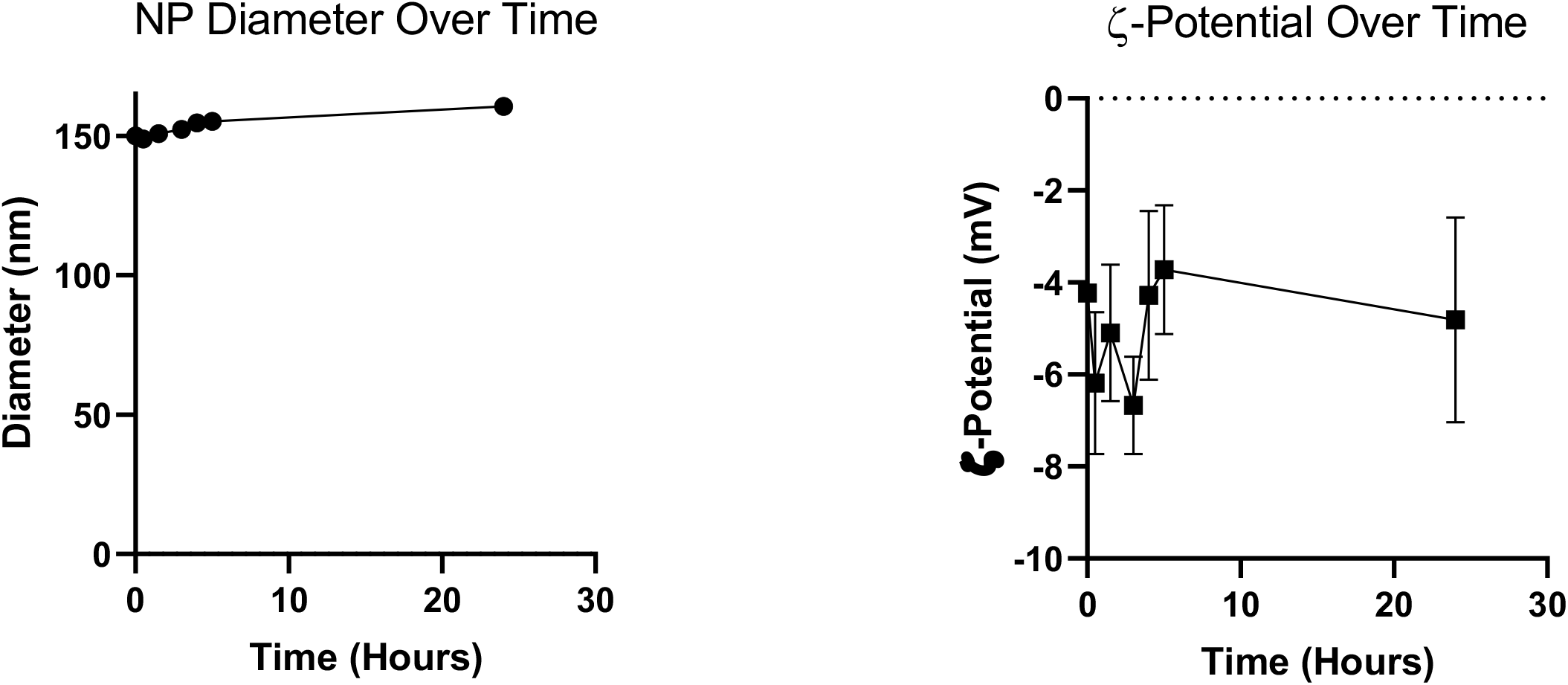
100 nm PSPEGRf/D=4.9 Stability Formulation Stability over Time in EGM-2. **A)** Dynamic Light Scattering (DLS) measurement of PEGylated NP diameter and **B)** Phase Analysis Light Scattering (PALS) measurement of NP ζ – potential over 24 hours. (n=5)

**Supplemental Image 3:**
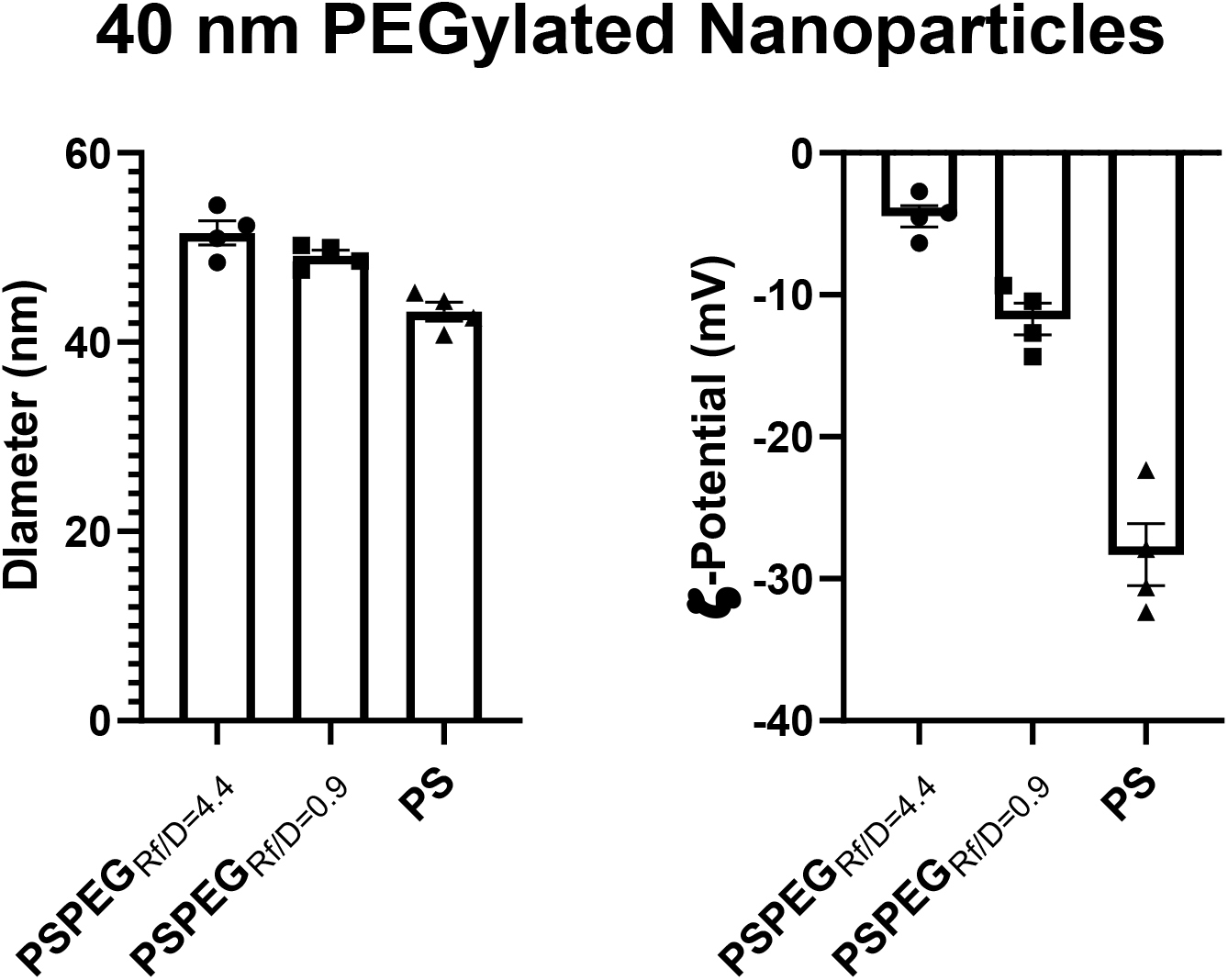
40 nm PSPEG Formulation: **A)** Dynamic Light Scattering (DLS) measurement of PEGylated NP diameter and **B)** Phase Analysis Light Scattering (PALS) measurement of NP ζ – potential. Data shown as mean ± SEM (n = 4)

**Supplemental Image 4:**
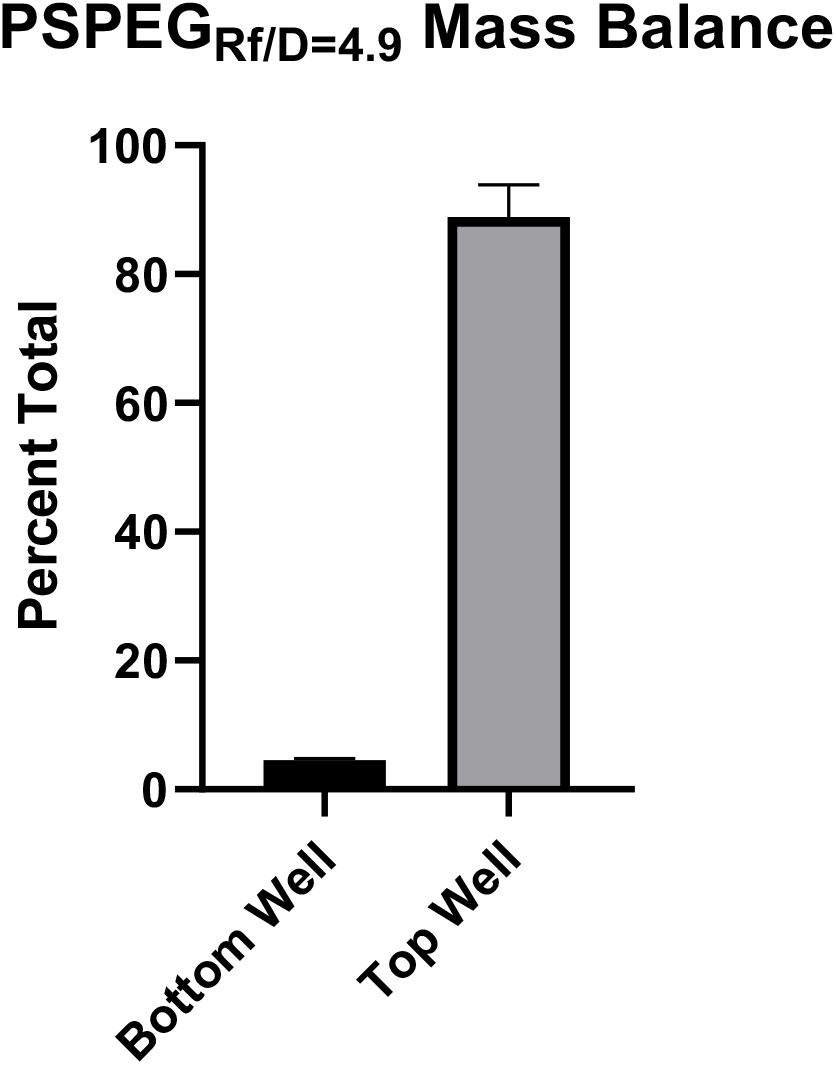
Percent of NP transported across LEC monolayer and in the top well after 24 hours for PSPEG_Rf/D=4.9_. (n=4)

**Supplemental Image 5:**
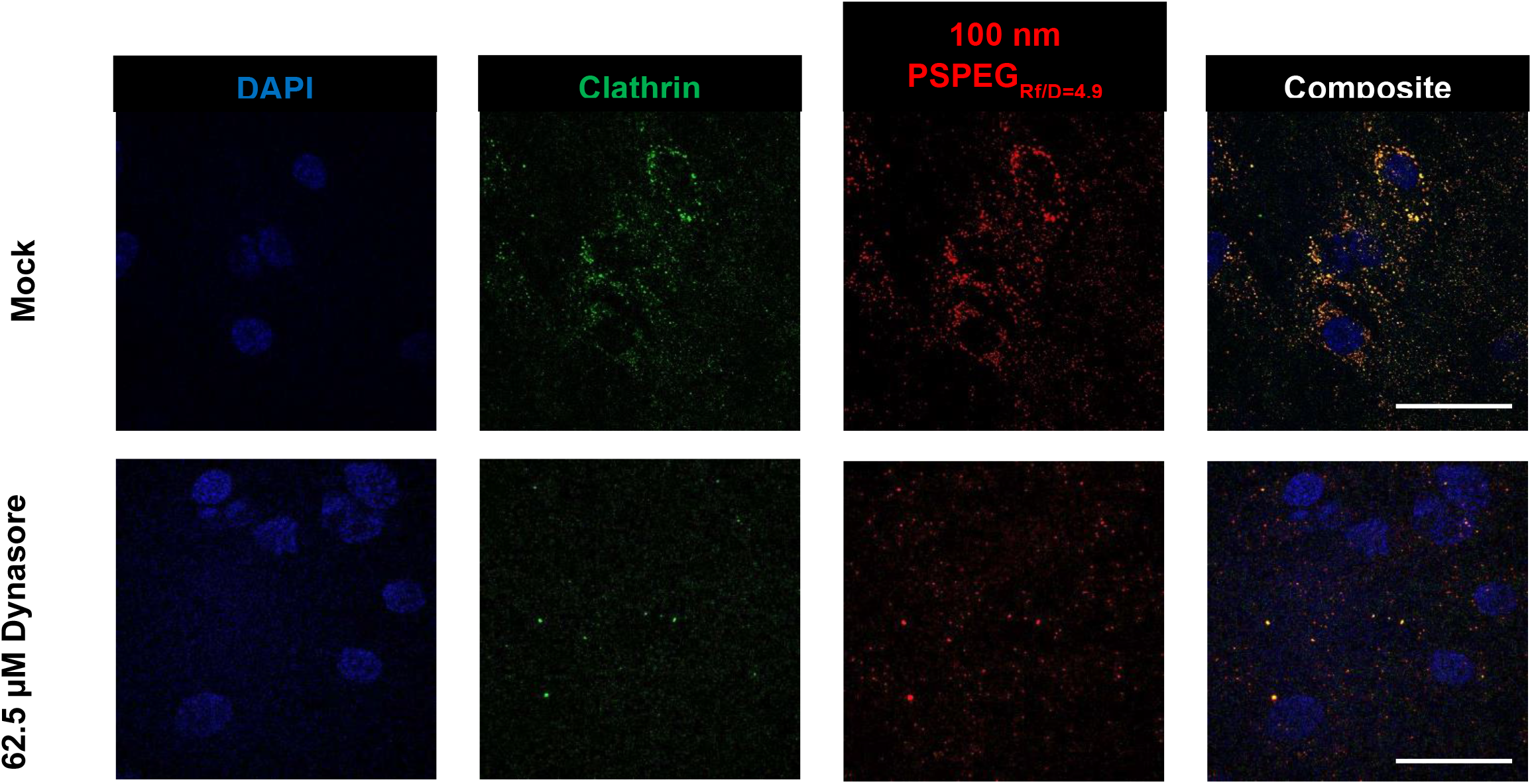
Confocal fluorescence image of PSPEG_Rf/D=4.9_ within LECs treated with the vehicle control and of PSPEG_Rf/D=4.9_ within LECs treated with Dynasore. Scale bar: 30 μm. (n = 3-4)

## References

[1] H. Wiig, M.A. Swartz, Interstitial fluid and lymph formation and transport: physiological regulation and roles in inflammation and cancer, Physiol Rev 92(3) (2012) 1005–60.

[2] S.T. Reddy, A. Rehor, H.G. Schmoekel, J.A. Hubbell, M.A. Swartz, In vivo targeting of dendritic cells in lymph nodes with poly(propylene sulfide) nanoparticles, Journal of Controlled Release 112(1) (2006) 26–34.

[3] V. Manolova, A. Flace, M. Bauer, K. Schwarz, P. Saudan, M.F. Bachmann, Nanoparticles target distinct dendritic cell populations according to their size, European Journal of Immunology 38(5) (2008) 1404–1413.

[4] H. Kobayashi, S. Kawamoto, M. Bernardo, M.W. Brechbiel, M.V. Knopp, P.L. Choyke, Delivery of gadolinium-labeled nanoparticles to the sentinel lymph node: comparison of the sentinel node visualization and estimations of intra-nodal gadolinium concentration by the magnetic resonance imaging, J Control Release 111(3) (2006) 343–351.

[5] E.M. Varypataki, A.L. Silva, C. Barnier-Quer, N. Collin, F. Ossendorp, W. Jiskoot, Synthetic long peptide-based vaccine formulations for induction of cell mediated immunity: A comparative study of cationic liposomes and PLGA nanoparticles, J Control Release 226 (2016) 98–106.

[6] S.M. Moghimi, A.E. Hawley, N.M. Christy, T. Gray, L. Illum, S.S. Davis, Surface engineered nanospheres with enhanced drainage into lymphatics and uptake by macrophages of the regional lymph nodes, FEBS Lett 344(1) (1994) 25–30.

[7] Q. Zeng, H. Jiang, T. Wang, Z. Zhang, T. Gong, X. Sun, Cationic micelle delivery of Trp2 peptide for efficient lymphatic draining and enhanced cytotoxic T-lymphocyte responses, Journal of Controlled Release 200 (2015) 1–12.

[8] S. De Koker, J. Cui, N. Vanparijs, L. Albertazzi, J. Grooten, F. Caruso, B.G. De Geest, Engineering Polymer Hydrogel Nanoparticles for Lymph Node-Targeted Delivery, Angewandte Chemie International Edition 55(4) (2016) 1334–1339.

[9] D.A. Rao, M.L. Forrest, A.W.G. Alani, G.S. Kwon, J.R. Robinson, Biodegradable PLGA based nanoparticles for sustained regional lymphatic drug delivery, J Pharm Sci 99(4) (2010) 2018–2031.

[10] L.M. Kaminskas, J. Kota, V.M. McLeod, B.D. Kelly, P. Karellas, C.J.H. Porter, PEGylation of polylysine dendrimers improves absorption and lymphatic targeting following SC administration in rats, Journal of Controlled Release 140(2) (2009) 108–116.

[11] S.M. Moghimi, A.C. Hunter, J.C. Murray, Long-circulating and target-specific nanoparticles: theory to practice, Pharmacol Rev 53(2) (2001) 283–318.

[12] M.E. Jung Soo Suk and Qingguo Xu and Namho Kim and Justin Hanes and Laura, PEGylation as a strategy for improving nanoparticle-based drug and gene delivery, Advanced Drug Delivery Reviews 99 (2016) 28–51.

[13] F. Alexis, E. Pridgen, L.K. Molnar, O.C. Farokhzad, Factors affecting the clearance and biodistribution of polymeric nanoparticles, Mol Pharm 5(4) (2008) 505–15.

[14] A. Gessner, R. Waicz, A. Lieske, B. Paulke, K. Mäder, R.H. Müller, Nanoparticles with decreasing surface hydrophobicities: influence on plasma protein adsorption, Int J Pharm 196(2) (2000) 245–9.

[15] C.D. Walkey, J.B. Olsen, H. Guo, A. Emili, W.C. Chan, Nanoparticle size and surface chemistry determine serum protein adsorption and macrophage uptake, J Am Chem Soc 134(4) (2012) 2139–47.

[16] H. Otsuka, Y. Nagasaki, K. Kataoka, PEGylated nanoparticles for biological and pharmaceutical applications, Advanced Drug Delivery Reviews 55(3) (2003) 403–419.

[17] J.V. Jokerst, T. Lobovkina, R.N. Zare, S.S. Gambhir, Nanoparticle PEGylation for imaging and therapy, Nanomedicine (Lond) 6(4) (2011) 715–728.

[18] Q. Xu, L.M. Ensign, N.J. Boylan, A. Schön, X. Gong, J.-C. Yang, N.W. Lamb, S. Cai, T. Yu, E. Freire, J. Hanes, Impact of Surface Polyethylene Glycol (PEG) Density on Biodegradable Nanoparticle Transport in Mucus ex Vivo and Distribution in Vivo, ACS Nano 9(9) (2015) 9217–9227.

[19] Q. Xu, N.J. Boylan, S. Cai, B. Miao, H. Patel, J. Hanes, Scalable method to produce biodegradable nanoparticles that rapidly penetrate human mucus, J Control Release 170(2) (2013) 279–86.

[20] E.A. Nance, G.F. Woodworth, K.A. Sailor, T.-Y. Shih, Q. Xu, G. Swaminathan, D. Xiang, C. Eberhart, J. Hanes, A dense poly(ethylene glycol) coating improves penetration of large polymeric nanoparticles within brain tissue, Sci Transl Med 4(149) (2012) 149ra119–149ra119.

[21] J.G. Dancy, A.S. Wadajkar, C.S. Schneider, J.R.H. Mauban, O.G. Goloubeva, G.F. Woodworth, J.A. Winkles, A.J. Kim, Non-specific binding and steric hindrance thresholds for penetration of particulate drug carriers within tumor tissue, J Control Release 238 (2016) 139–148.

[22] V. Triacca, E. Güç, W.W. Kilarski, M. Pisano, M.A. Swartz, Transcellular Pathways in Lymphatic Endothelial Cells Regulate Changes in Solute Transport by Fluid Stress, Circ Res 120(9) (2017) 1440–1452.

[23] W.J. Kelley, C.A. Fromen, G. Lopez-Cazares, O. Eniola-Adefeso, PEGylation of model drug carriers enhances phagocytosis by primary human neutrophils, Acta Biomater 79 (2018) 283–293.

[24] K. Shameli, M. Bin Ahmad, S.D. Jazayeri, S. Sedaghat, P. Shabanzadeh, H. Jahangirian, M. Mahdavi, Y. Abdollahi, Synthesis and Characterization of Polyethylene Glycol Mediated Silver Nanoparticles by the Green Method, International Journal of Molecular Sciences 13(6) (2012).

[25] H. Wu, S. Nagarajan, L. Zhou, Y. Duan, J. Zhang, Synthesis and characterization of cellulose nanocrystal-graft-poly(d-lactide) and its nanocomposite with poly(l-lactide), Polymer 103 (2016) 365–375.

[26] S. Hirosue, E. Vokali, V.R. Raghavan, M. Rincon-Restrepo, A.W. Lund, P. Corthésy-Henrioud, F. Capotosti, C. Halin Winter, S. Hugues, M.A. Swartz, Steady-state antigen scavenging, cross-presentation, and CD8+ T cell priming: a new role for lymphatic endothelial cells, J Immunol 192(11) (2014) 5002–11.

[27] Q. Xu, L.M. Ensign, N.J. Boylan, A. Schön, X. Gong, J.C. Yang, N.W. Lamb, S. Cai, T. Yu, E. Freire, J. Hanes, Impact of Surface Polyethylene Glycol (PEG) Density on Biodegradable Nanoparticle Transport in Mucus ex Vivo and Distribution in Vivo, ACS Nano 9(9) (2015) 9217–27.

[28] P. Guo, J. Yang, D. Jia, M.A. Moses, D.T. Auguste, ICAM-1-Targeted, Lcn2 siRNA-Encapsulating Liposomes are Potent Anti-angiogenic Agents for Triple Negative Breast Cancer, Theranostics 6(1) (2016) 1–13.

[29] E. Sari, Y. Tunc-Sarisozen, H. Mutlu, R. Shahbazi, G. Ucar, K. Ulubayram, ICAM-1 targeted catalase encapsulated PLGA-b-PEG nanoparticles against vascular oxidative stress, J Microencapsul 32(7) (2015) 687–98.

[30] L. Del Amo, A. Cano, M. Ettcheto, E.B. Souto, M. Espina, A. Camins, M.L. García, E. Sánchez-López, Surface Functionalization of PLGA Nanoparticles to Increase Transport across the BBB for Alzheimer’s Disease, Applied Sciences 11(9) (2021).

[31] Q. Lin, P. Fathi, X. Chen, Nanoparticle delivery <em>in vivo</em>: A fresh look from intravital imaging, EBioMedicine 59 (2020).

[32] S.F. Tehrani, F. Bernard-Patrzynski, I. Puscas, G. Leclair, P. Hildgen, V.G. Roullin, Length of surface PEG modulates nanocarrier transcytosis across brain vascular endothelial cells, Nanomedicine: Nanotechnology, Biology and Medicine 16 (2019) 185–194.

[33] J.-M. Rabanel, P.-A. Piec, S. Landri, S.A. Patten, C. Ramassamy, Transport of PEGylated-PLA nanoparticles across a blood brain barrier model, entry into neuronal cells and in vivo brain bioavailability, Journal of Controlled Release 328 (2020) 679–695.

[34] M. Kim, M. Jeong, S. Hur, Y. Cho, J. Park, H. Jung, Y. Seo, H.A. Woo, K.T. Nam, K. Lee, H. Lee, Engineered ionizable lipid nanoparticles for targeted delivery of RNA therapeutics into different types of cells in the liver, Sci Adv 7(9) (2021).

[35] R.M. Williams, J. Shah, B.D. Ng, D.R. Minton, L.J. Gudas, C.Y. Park, D.A. Heller, Mesoscale Nanoparticles Selectively Target the Renal Proximal Tubule Epithelium, Nano Letters 15(4) (2015) 2358–2364.

[36] H. Zhou, Z. Fan, P.Y. Li, J. Deng, D.C. Arhontoulis, C.Y. Li, W.B. Bowne, H. Cheng, Dense and Dynamic Polyethylene Glycol Shells Cloak Nanoparticles from Uptake by Liver Endothelial Cells for Long Blood Circulation, ACS Nano 12(10) (2018) 10130–10141.

[37] X.J. Du, J.L. Wang, W.W. Liu, J.X. Yang, C.Y. Sun, R. Sun, H.J. Li, S. Shen, Y.L. Luo, X.D. Ye, Y.H. Zhu, X.Z. Yang, J. Wang, Regulating the surface poly(ethylene glycol) density of polymeric nanoparticles and evaluating its role in drug delivery in vivo, Biomaterials 69 (2015) 1–11.

[38] Y. Li, M. Kröger, W.K. Liu, Endocytosis of PEGylated nanoparticles accompanied by structural and free energy changes of the grafted polyethylene glycol, Biomaterials 35(30) (2014) 8467–8478.

[39] M. Li, S. Jiang, J. Simon, D. Paßlick, M.-L. Frey, M. Wagner, V. Mailänder, D. Crespy, K. Landfester, Brush Conformation of Polyethylene Glycol Determines the Stealth Effect of Nanocarriers in the Low Protein Adsorption Regime, Nano Letters 21(4) (2021) 1591–1598.

[40] J.L. Perry, K.G. Reuter, M.P. Kai, K.P. Herlihy, S.W. Jones, J.C. Luft, M. Napier, J.E. Bear, J.M. DeSimone, PEGylated PRINT nanoparticles: the impact of PEG density on protein binding, macrophage association, biodistribution, and pharmacokinetics, Nano Lett 12(10) (2012) 5304–10.

[41] Q. Yang, S.W. Jones, C.L. Parker, W.C. Zamboni, J.E. Bear, S.K. Lai, Evading Immune Cell Uptake and Clearance Requires PEG Grafting at Densities Substantially Exceeding the Minimum for Brush Conformation, Molecular Pharmaceutics 11(4) (2014) 1250–1258.

[42] Z. Ait Bachir, Y. Huang, M. He, L. Huang, X. Hou, R. Chen, F. Gao, Effects of PEG surface density and chain length on the pharmacokinetics and biodistribution of methotrexate-loaded chitosan nanoparticles, Int J Nanomedicine 13 (2018) 5657–5671.

[43] D. Pozzi, V. Colapicchioni, G. Caracciolo, S. Piovesana, A.L. Capriotti, S. Palchetti, S. De Grossi, A. Riccioli, H. Amenitsch, A. Laganà, Effect of polyethyleneglycol (PEG) chain length on the bio-nano-interactions between PEGylated lipid nanoparticles and biological fluids: from nanostructure to uptake in cancer cells, Nanoscale 6(5) (2014) 2782–92.

[44] L.B. Sims, L.T. Curtis, H.B. Frieboes, J.M. Steinbach-Rankins, Enhanced uptake and transport of PLGA-modified nanoparticles in cervical cancer, J Nanobiotechnology 14 (2016) 33–33.

[45] G. Aguilera, C.C. Berry, R.M. West, E. Gonzalez-Monterrubio, A. Angulo-Molina, Ó. Arias-Carrión, M.Á. Méndez-Rojas, Carboxymethyl cellulose coated magnetic nanoparticles transport across a human lung microvascular endothelial cell model of the blood–brain barrier, Nanoscale Advances 1(2) (2019) 671–685.

[46] S.I. Ahn, Y.J. Sei, H.-J. Park, J. Kim, Y. Ryu, J.J. Choi, H.-J. Sung, T.J. MacDonald, A.I. Levey, Y. Kim, Microengineered human blood–brain barrier platform for understanding nanoparticle transport mechanisms, Nature Communications 11(1) (2020) 175.

[47] N. Chattopadhyay, J. Zastre, H.L. Wong, X.Y. Wu, R. Bendayan, Solid lipid nanoparticles enhance the delivery of the HIV protease inhibitor, atazanavir, by a human brain endothelial cell line, Pharm Res 25(10) (2008) 2262–71.

[48] V. Francia, A. Aliyandi, A. Salvati, Effect of the development of a cell barrier on nanoparticle uptake in endothelial cells, Nanoscale 10(35) (2018) 16645–16656.

[49] H.R. Kim, S. Gil, K. Andrieux, V. Nicolas, M. Appel, H. Chacun, D. Desmaële, F. Taran, D. Georgin, P. Couvreur, Low-density lipoprotein receptor-mediated endocytosis of PEGylated nanoparticles in rat brain endothelial cells, Cell Mol Life Sci 64(3) (2007) 356–64.

[50] S.M. Moghimi, D. Simberg, Nanoparticle transport pathways into tumors, Journal of Nanoparticle Research 20(6) (2018) 169.

[51] T. Skotland, K. Sandvig, Transport of nanoparticles across the endothelial cell layer, Nano Today 36 (2021) 101029.

[52] J. Kreuter, Mechanism of polymeric nanoparticle-based drug transport across the blood-brain barrier (BBB), J Microencapsul 30(1) (2013) 49–54.

[53] Z. Wang, C. Tiruppathi, R.D. Minshall, A.B. Malik, Size and Dynamics of Caveolae Studied Using Nanoparticles in Living Endothelial Cells, ACS Nano 3(12) (2009) 4110–4116.

[54] Z. Wang, C. Tiruppathi, J. Cho, R.D. Minshall, A.B. Malik, Delivery of nanoparticle: complexed drugs across the vascular endothelial barrier via caveolae, IUBMB Life 63(8) (2011) 659–67.

[55] D. Ye, M.N. Raghnaill, M. Bramini, E. Mahon, C. Åberg, A. Salvati, K.A. Dawson, Nanoparticle accumulation and transcytosis in brain endothelial cell layers, Nanoscale 5(22) (2013) 11153–11165.

[56] R. Roozendaal, R.E. Mebius, G. Kraal.

[57] R.G. Thorne, C. Nicholson, <em>In vivo</em> diffusion analysis with quantum dots and dextrans predicts the width of brain extracellular space, Proceedings of the National Academy of Sciences 103(14) (2006) 5567.

[58] T. Stylianopoulos, M.-Z. Poh, N. Insin, M.G. Bawendi, D. Fukumura, L.L. Munn, R.K. Jain, Diffusion of particles in the extracellular matrix: the effect of repulsive electrostatic interactions, Biophysical journal 99(5) (2010) 1342–1349.

[59] O. Lieleg, R.M. Baumgärtel, A.R. Bausch, Selective filtering of particles by the extracellular matrix: an electrostatic bandpass, Biophysical journal 97(6) (2009) 1569–1577.

[60] M. Le Goas, F. Testard, O. Taché, N. Debou, B. Cambien, G. Carrot, J.-P. Renault, How Do Surface Properties of Nanoparticles Influence Their Diffusion in the Extracellular Matrix? A Model Study in Matrigel Using Polymer-Grafted Nanoparticles, Langmuir 36(35) (2020) 10460–10470.

[61] J. Hansing, C. Ciemer, W.K. Kim, X. Zhang, J.E. DeRouchey, R.R. Netz, Nanoparticle filtering in charged hydrogels: Effects of particle size, charge asymmetry and salt concentration, The European Physical Journal E 39(5) (2016) 53.

[62] B. Mattix, T. Moore, O. Uvarov, S. Pollard, L. O’Donnell, K. Park, D. Horne, J. Dhulekar, L. Reese, D. Nguyen, J. Kraveka, K. Burg, F. Alexis, EFFECTS OF POLYMERIC NANOPARTICLE SURFACE PROPERTIES ON INTERACTION WITH BRAIN TUMOR ENVIRONMENT, Nano LIFE 3(4) (2013) 1343003–1343003.

[63] L. Tomasetti, R. Liebl, D.S. Wastl, M. Breunig, Influence of PEGylation on nanoparticle mobility in different models of the extracellular matrix, Eur J Pharm Biopharm 108 (2016) 145–155.

[64] H.I. Labouta, M.J. Gomez-Garcia, C.D. Sarsons, T. Nguyen, J. Kennard, W. Ngo, K. Terefe, N. Iragorri, P. Lai, K.D. Rinker, D.T. Cramb, Surface-grafted polyethylene glycol conformation impacts the transport of PEG-functionalized liposomes through a tumour extracellular matrix model, RSC Advances 8(14) (2018) 7697–7708.

[65] S.K. Lai, Y.-Y. Wang, J. Hanes, Mucus-penetrating nanoparticles for drug and gene delivery to mucosal tissues, Advanced Drug Delivery Reviews 61(2) (2009) 158–171.

